# Survival-informed mutational network clustering reveals prognostic glioma architectures

**DOI:** 10.64898/2026.11.12.766165

**Authors:** Beatriz N. Leitão, Jack Kuipers, Niko Beerenwinkel, Susana Vinga

## Abstract

Cancer heterogeneity remains a major challenge for predicting patient outcomes. To address this challenge, we introduce SNOWS (Survival-iNfOrmed netWork cluStering), a probabilistic framework that integrates somatic mutations, clinical covariates and survival information to discover patient groups with distinct mutation patterns and survival profiles, using Bayesian networks to capture dependencies involving mutations and clinical variables. Applied to 950 adult-type diffuse gliomas from TCGA, SNOWS identified three clusters that transferred successfully to classify an independent CGGA cohort of 181 patients using mutation and clinical data, without requiring their survival outcomes. The transferred clusters showed distinct survival, mutation profiles and dependency structures, and provided prognostic information beyond WHO 2021 tumour subtype, age and sex, significantly increasing the C-index from 0.691 to 0.712. Cluster assignments were driven by specific mutation frequencies, with network dependencies improving the assignments of patients with less typical molecular profiles. Beyond glioma, SNOWS also improved prognostic discrimination in pan-kidney and pan-myeloid cohorts. These results support survival-informed network clustering as a general framework for resolving clinically relevant cancer heterogeneity.

## 1 Introduction

Cancer heterogeneity remains a major challenge for prognostic prediction [1]. Large-scale whole-genome sequencing initiatives have enabled the characterisation of somatic mutational landscapes across cancer types, revealing patterns of genomic alteration that contribute to tumour heterogeneity [2]. These mutational profiles provide a rich basis for computational approaches aimed at identifying molecularly distinct patient groups. A challenge in precision oncology is therefore to integrate and interpret such high-dimensional data in order to stratify patients into clinically meaningful subgroups [3]. Such stratification may support more accurate diagnosis, prognosis, and risk assessment, and ultimately inform treatment decision-making.

Gliomas provide a particularly relevant setting in which to investigate this challenge. These primary brain tumours are thought to arise from neural stem or progenitor cells carrying tumour-initiating genetic alterations [4]. According to the 2021 World Health Organisation (WHO) classification of central nervous system (CNS) tumours, gliomas comprise a heterogeneous group of entities defined by histological and molecular features [5]. The broad category of glioma accounts for approximately 24% of all primary brain and other CNS tumours and 81% of malignant brain tumours, underscoring their major clinical relevance [6]. Adult-type diffuse gliomas represent more than 90% of gliomas and include three biologically and prognostically distinct tumour types: isocitrate dehydrogenase (IDH)-mutant astrocytoma (ASTRO), IDH-mutant and 1p/19q-codeleted oligodendroglioma (OLIGO), and IDH-wild-type glioblastoma (GBM) [4].

Although adult-type diffuse gliomas are generally considered incurable, their prognosis is highly heterogeneous. In a national-level US cohort, 1-year overall survival (OS) was 53.7% for WHO grade 4 glioblastoma, compared with 98.0%, 92.4%, and 76.3% for WHO grade 2, 3, and 4 astrocytoma, respectively, and 97.9% and 94.4% for WHO grade 2 and 3 oligodendroglioma, respectively [7]. The current WHO 2021 classification therefore captures a major component of glioma biology and prognosis.

However, recent studies have revealed additional molecular heterogeneity within the major WHO 2021 adult-type diffuse glioma entities, enabling the identification of prognostically distinct molecular groups within astrocytoma [8, 9] and oligodendroglioma [10]. In glioblastoma, transcriptomic profiling combined with machine-learning feature selection and regularised Cox regression has identified multigene signatures capable of stratifying patients by survival [11]. For an independent Chinese Glioma Genome Atlas (CGGA) cohort [12], the prognostic separation could also be reproduced [11]. Across the three major adult-type diffuse glioma entities and a group titled ‘other glioma’ that combined rarer entities, routinely collected mutational and other molecular alterations have also been used to identify patient groups that provided modestly better survival prediction than the WHO 2021 classification [13]. Furthermore, Ruffle et al. [13] inferred a cohort-wide network of statistical relationships among these mutational and other molecular alterations, suggesting that while each of these features has some prognostic value on its own, their interactions provide additional insight. This observation motivates approaches that explicitly model relationships among molecular alterations.

Network-based patient clustering can capture molecular relationships overlooked when mutations are considered independently, either by incorporating prior gene-interaction information [14] or learning cluster-specific dependencies from tumour data [15, 16]. However, these approaches do not use survival to guide cluster estimation, potentially overlooking prognostically relevant molecular structure.

Here, we develop SNOWS (**S**urvival-i**N**f**O**rmed net**W**ork clu**S**tering), a probabilistic framework that identifies patient clusters through cluster-specific Bayesian networks (BNs) capturing conditional dependencies among somatic mutations and clinical covariates. Survival information guides the discovery of network-defined clusters with distinct prognostic profiles, while established clinical sources of variation are accounted for in both the network and survival components. Once trained, the model assigns previously unseen patients using mutational and clinical data alone, without requiring survival outcomes.

We applied SNOWS to adult-type diffuse glioma, using The Cancer Genome Atlas (TCGA) for discovery and an independent CGGA cohort for validation. SNOWS identified three externally reproducible patient clusters with distinct survival outcomes, mutational profiles, and dependency architectures. These clusters provided prognostic information beyond WHO 2021 tumour subtype and clinical covariates and improved survival prediction relative to alternative network-based and conventional clustering approaches. We further evaluated SNOWS generalisability in additional cancer cohorts, namely TCGA pan-kidney cancer and MD Anderson pan-myeloid malignancy cohorts. Together, these analyses support survival-informed network clustering for refining patient stratification and uncovering prognostically relevant molecular architectures.

## 2 Methods

### 2.1 Model formulation

We developed SNOWS, a probabilistic clustering framework in which patient subtypes are defined jointly by a BN that integrates mutations and clinical variables together with subtype-specific survival behaviour (Fig. 1). SNOWS extends CANclust [16] by incorporating survival information into BN clustering. Cluster-specific BNs model dependencies among mutations and clinical covariates, while survival information guides the identification of prognostically distinct clusters. Clinical covariates enter both the BNs and survival models to account for clinical sources of variation, reducing the extent to which the resulting clusters are explained by these known clinical factors, facilitating the identification of additional molecular structure associated with prognosis. Once trained, SNOWS assigns new patients using only mutation and clinical data, without requiring survival outcomes.

**Figure 1:**
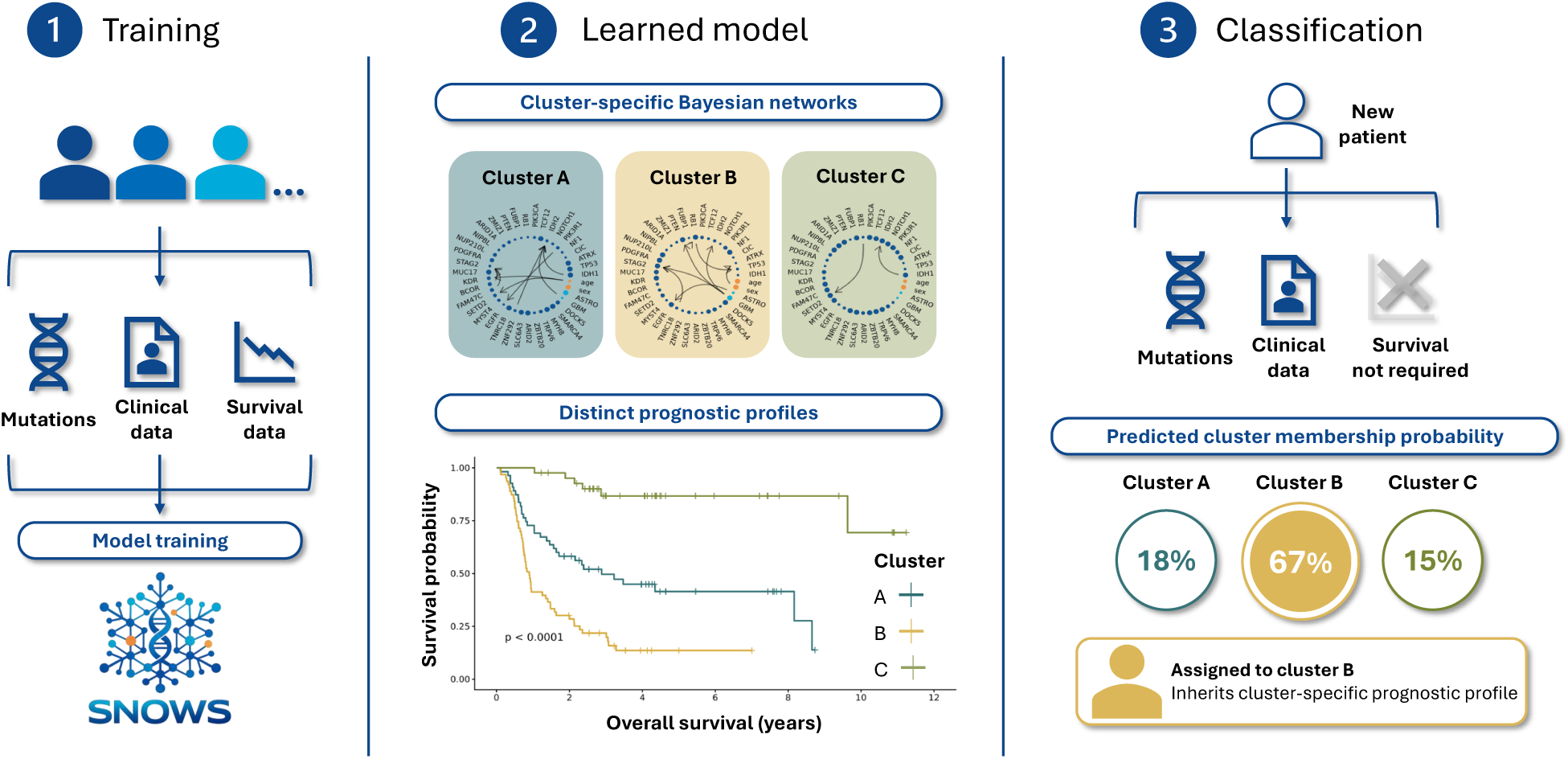
Workflow overview. During model training, somatic mutation data and clinical covariates define cluster-specific Bayesian networks, while survival information guides discovery of clusters with different survival profiles. The fitted model is subsequently applied to a new patient using mutation and clinical data only; survival time and event status are not used for assignment. The resulting cluster assignment can then be used for downstream prognostic stratification.

Patients are indexed by *i* = 1, … , *N* and latent clusters by *k* = 1, … , *K*, with *V* denoting the set of mutational variables and *C* the set of clinical covariates. For each patient *i*, we observe a mutational profile *X_V__iᵢ_*, a set of clinical covariates *X_C__iᵢ_*, and a survival outcome (*t_i_, δ_i_*), where *t_i_ >* 0 denotes the observed follow-up time and *δ_i_ ∈* {0, 1} indicates whether the event was observed. Latent cluster assignment is represented by *Z_i_ ∈* {1, … , *K*}, with cluster-membership prior probabilities ***τ*** = (*τ*_1_, … , *τ_K_*) satisfying 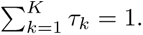

For each cluster *k*, SNOWS defines a BN with graph structure *G_k_* and parameters *θ_k_*. Survival outcomes are modelled through a parametric survival model with global regression coefficients ***β*** for clinical covariates, cluster-specific risk effects *γ_k_*, and a shape parameter *ν* shared across clusters. In both models, clinical covariates are included so that cluster-specific mutational dependencies and survival effects are estimated while accounting for established clinical sources of variation.

### 2.2 Bayesian network modelling

The BNs model cluster-specific dependencies among molecular mutations while accounting for clinical covariates. In each graph *G_k_*, nodes represent mutations and clinical covariates, while directed edges specify conditional probability distributions parametrised by *θ_k_*. Clinical covariates are restricted to outgoing edges, allowing their effects on the mutations variables to be accounted for without differences in these covariates directly driving cluster assignment. Mutation variables, in contrast, may have both incoming and outgoing edges, allowing their dependencies with other variables to be modelled [15, 16].

Under this formulation, the mutational profile of patient *i* is *X_V__i_* = (*X_ij_*)*_j∈V_* , where *X_ij_ ∈* {0, 1} denotes the mutation status of gene *j* for patient *i*. The mutational profile is modelled conditional on the clinical covariates *X_C__i_* . For cluster *k*,

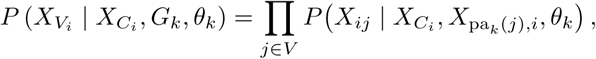

where pa*_k_* (*j*) denotes the parent set of mutational variable *j* in graph *G_k_*. The corresponding BN log-likelihood quantifies how well the patient’s mutational profile is explained by cluster *k*:

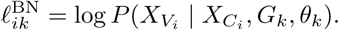

### 2.3 Weibull survival model and parameter estimation

SNOWS models survival time *T* using a Weibull distribution, *T ∼* Weibull(*λ, ν*), where *λ >* 0 is the scale parameter and *ν >* 0 is the shape parameter shared across clusters. Under this parameterisation, the hazard and survival functions are

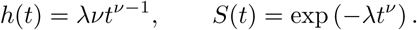

Clinical covariates are incorporated through a log-linear model for the hazard scaling parameter. During model fitting, uncertainty in cluster membership is represented through soft cluster-membership probabilities. Specifically, let *P_i_* = (*P_i_*_1_, … , *P_iK_*) denote the current membership probabilities for patient *i*. The scale parameter for patient *i* is defined as

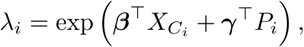

where ***β*** represents global regression coefficients for clinical covariates and ***γ*** = (*γ*_1_, … , *γ_K_*) are cluster-specific survival coefficients. This formulation ensures that all patients contribute to the estimation of cluster-specific survival parameters in proportion to their uncertainty, rather than being assigned deterministically to a single cluster. When evaluating the survival likelihood of patient *i* under a specific cluster *k*, cluster membership is treated as fixed. In this case, the hazard scaling parameter is defined as

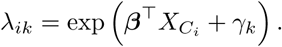

This can be regarded as evaluating the survival likelihood under a one-hot cluster assignment. The individual survival log-likelihood contribution [17] under cluster *k* is therefore

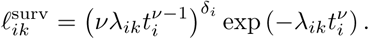

### 2.4 Integrated clustering

The BN and survival log-likelihood contributions arise from different models and are therefore not on the same numerical scale. To balance their contributions to clustering, we apply global exponential-mean normalisation to each component before combining. Let 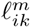 denote the unnormalised log-likelihood contribution of modality *m ∈* {BN, surv} for patient *i* under cluster *k*. The normalised log-likelihood is defined as

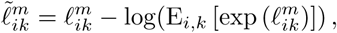

where the expectation is taken across all patients and clusters, ensuring that 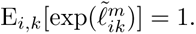

The normalised components are then combined into a cluster-specific score:

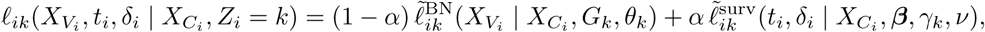

where *α ∈* [0, 1] controls the relative weighting of the BN and survival information. Normalisation and *α* play distinct roles: the former is applied separately to each log-likelihood before combination, whereas the latter determines their relative weights. Consequently, normalisation cannot be absorbed into *α*. When *α* = 0, SNOWS reduces to CANclust [16], with clustering driven entirely by the BN component. When *α* = 1, clustering depends solely on survival information, conceptually resembling previous survival-based clustering approaches [18]. Intermediate values allow both sources of information to jointly guide cluster discovery.

SNOWS is fitted using an expectation-maximisation (EM) algorithm that alternates between model parameter estimation and posterior cluster-membership updates. The log-likelihood objective is

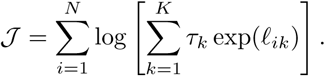

At each iteration, BN structures and parameters and the survival parameters are updated conditional on the current cluster-membership probabilities. Cluster priors are estimated using Dirichlet smoothing:

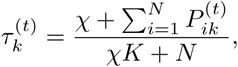

where *χ >* 0 is a pseudocount. Posterior cluster-membership probabilities are then updated as follows:

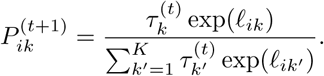

The complete fitting procedure is summarised in Algorithm 1.

#### Algorithm 1: SNOWS

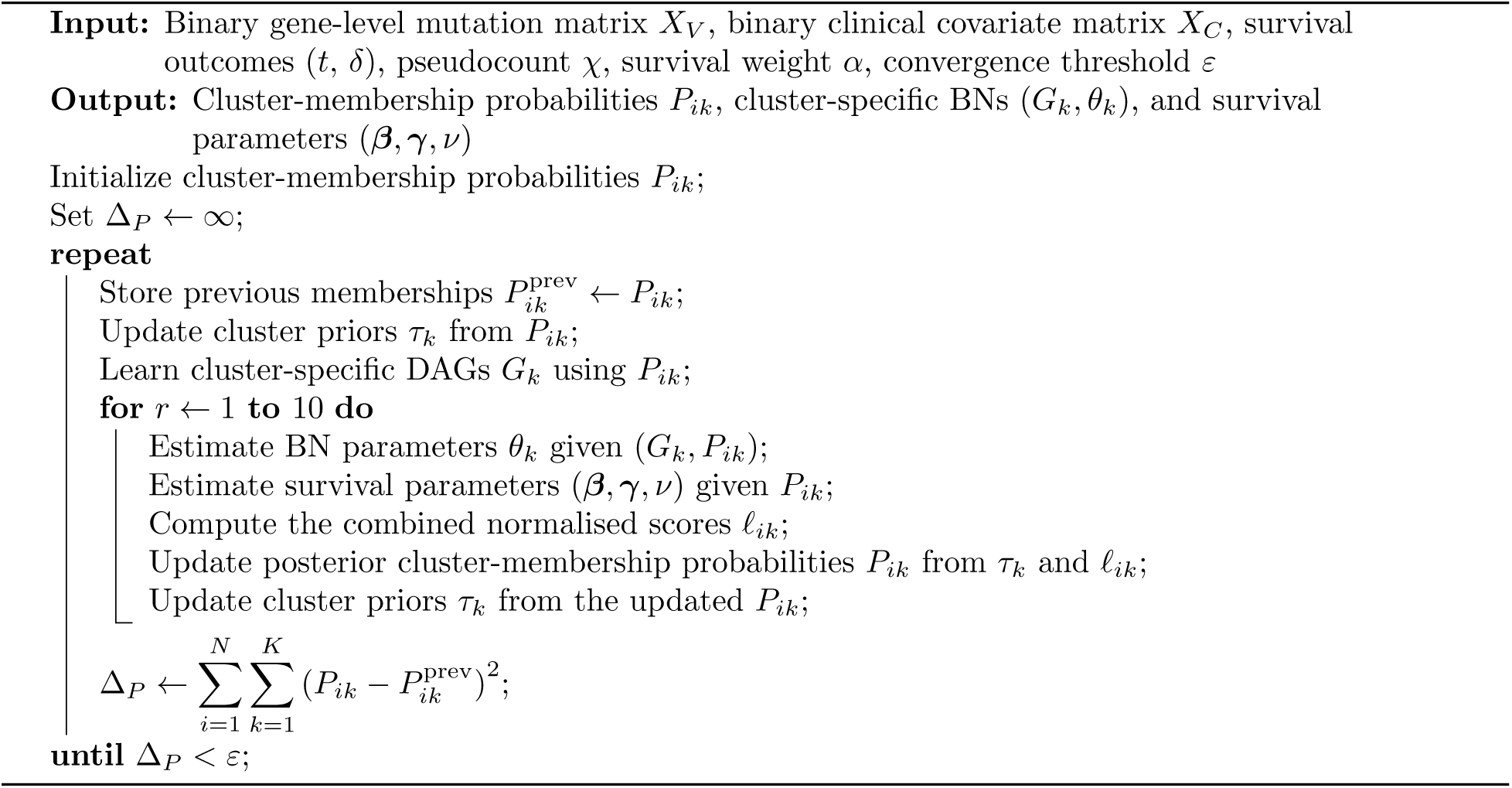

After model training, new patients are assigned to clusters using only the fitted BN models and cluster priors, without survival outcomes. Given mutation and clinical data (*X_V__i_ , X_C__i_* ), posterior membership probabilities are computed as

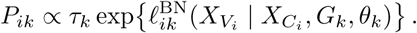

These probabilities define soft cluster memberships. For analyses requiring discrete assignments, each patient is assigned to the cluster with the highest posterior probability, *Z*^^^*_i_* = arg max*_k_ P_ik_*.

### 2.5 Data

Four cancer cohorts (Supplementary Section A.1) were analysed: TCGA glioma (GBMLGG), an independent CGGA glioma cohort, TCGA pan-kidney (KIPAN), and a pan-myeloid malignancy cohort from MD Anderson Cancer Center.

Somatic mutation data for TCGA glioma and pan-kidney were obtained from FireBrowse using GDAC Firehose Mutation Analysis outputs generated with MutSig v2.0 [19]. Mutation Annotation Format (MAF) files were converted into binary gene-level matrices indicating the presence of non-silent mutations. Genes were retained if significantly mutated (Benjamini–Hochberg false discovery rate *q ≤* 0.1) and present in at least 1% of patients, yielding 35 genes for glioma and 31 for pan-kidney. The glioma discovery cohort comprised 950 TCGA patients, reclassified according to WHO 2021 [20]. External validation used 181 patients from the CGGA cohort [12], similarly classified using available molecular and histological data [21]. CGGA mutation profiles were restricted to the 35 TCGA-selected genes to ensure consistent features across cohorts. The pan-kidney cohort comprised 898 patients.

The pan-myeloid cohort comprised 1320 patients. Mutations were obtained from an 81-gene targeted sequencing panel [22] and encoded as binary gene-level indicators. After excluding features with prevalence below 1%, the dataset included 38 mutation variables, eight cytogenetic profiles, six blood-derived variables, and five bone-marrow-derived variables.

Across cohorts, clinical data included age, biological sex, disease subtype, OS time, and event status. Somatic mutations and cytogenetic variables were binarised, while continuous variables were dichotomised using median-based cutoffs. Detailed cohort characteristics are provided in Supplementary Table 1.

### 2.6 Evaluation metrics and model selection

The prognostic value of cluster assignment was assessed using the Cox log-likelihood gain, likelihood-ratio tests, Harrell’s concordance index (C-index), and integrated Brier score (IBS). Associations between cluster assignments and clinical variables were evaluated using normalised mutual information (NMI) and McFadden’s pseudo-*R*^2^. Detailed definitions and statistical procedures are provided in Supplementary Section A.2.

The number of clusters was selected using the Bayesian information criterion (BIC) on an edge-free BN mixture model (equivalent to a Bernoulli mixture model), evaluated across candidate cluster numbers and pseudocount values without survival information. For glioma, the BN pseudocount *χ* and survival weight *α* were selected using five-fold cross-validation (CV) within the TCGA discovery cohort, considering incremental prognostic performance and preservation of non-collapsed clusters. Parameter grids and sensitivity analyses are provided in the Supplementary Section A.3. For the pan-kidney and pan-myeloid cohorts, *χ* was fixed to the glioma-selected value (*χ* = 0.001), while *α* was selected by inner CV within each outer training fold, primarily based on mean ΔC-index and preservation of the selected number of clusters. Models were then refitted to each outer training set and evaluated on the corresponding held-out fold. Unless otherwise specified, implementation settings inherited from CANclust [16] were retained.

## 3 Results

### 3.1 External validation reveals added prognostic value

The WHO 2021 classification captures prognostic differences among adult-type diffuse gliomas, as reflected by distinct survival distributions in CGGA (Supplementary Fig. 6; global log-rank *p <* 0.0001). However, substantial heterogeneity remains within these tumour types. We therefore investigated whether SNOWS could identify additional prognostic structure after accounting for WHO subtype, age, and biological sex.

SNOWS was trained on TCGA glioma (*n* = 950) and externally validated in CGGA (*n* = 181). The TCGA-derived BN structures, parameters, and cluster priors were fixed and applied to CGGA, where patients were assigned using mutation and clinical data alone, without survival outcomes.

Model selection supported three clusters across the evaluated pseudocount values. Five-fold CV within TCGA selected *χ* = 0.001 and *α* = 0.04, corresponding to a model in which survival information contributed to cluster discovery while preserving all three clusters (Supplementary Fig. 5). The selected configuration was refitted to the complete TCGA cohort, yielding a training-set C-index of 0.784 and IBS of 0.106.

Applied to CGGA, SNOWS assigned 62, 75, and 44 patients to Clusters A, B, and C, respectively. OS differed significantly across clusters (global log-rank *p <* 0.0001; Fig. 2a), with the poorest survival in Cluster B, intermediate survival in Cluster A, and the most favourable in Cluster C. As CGGA survival outcomes were not used for assignment, this separation provides independent evidence of prognostic relevance.

**Figure 2:**
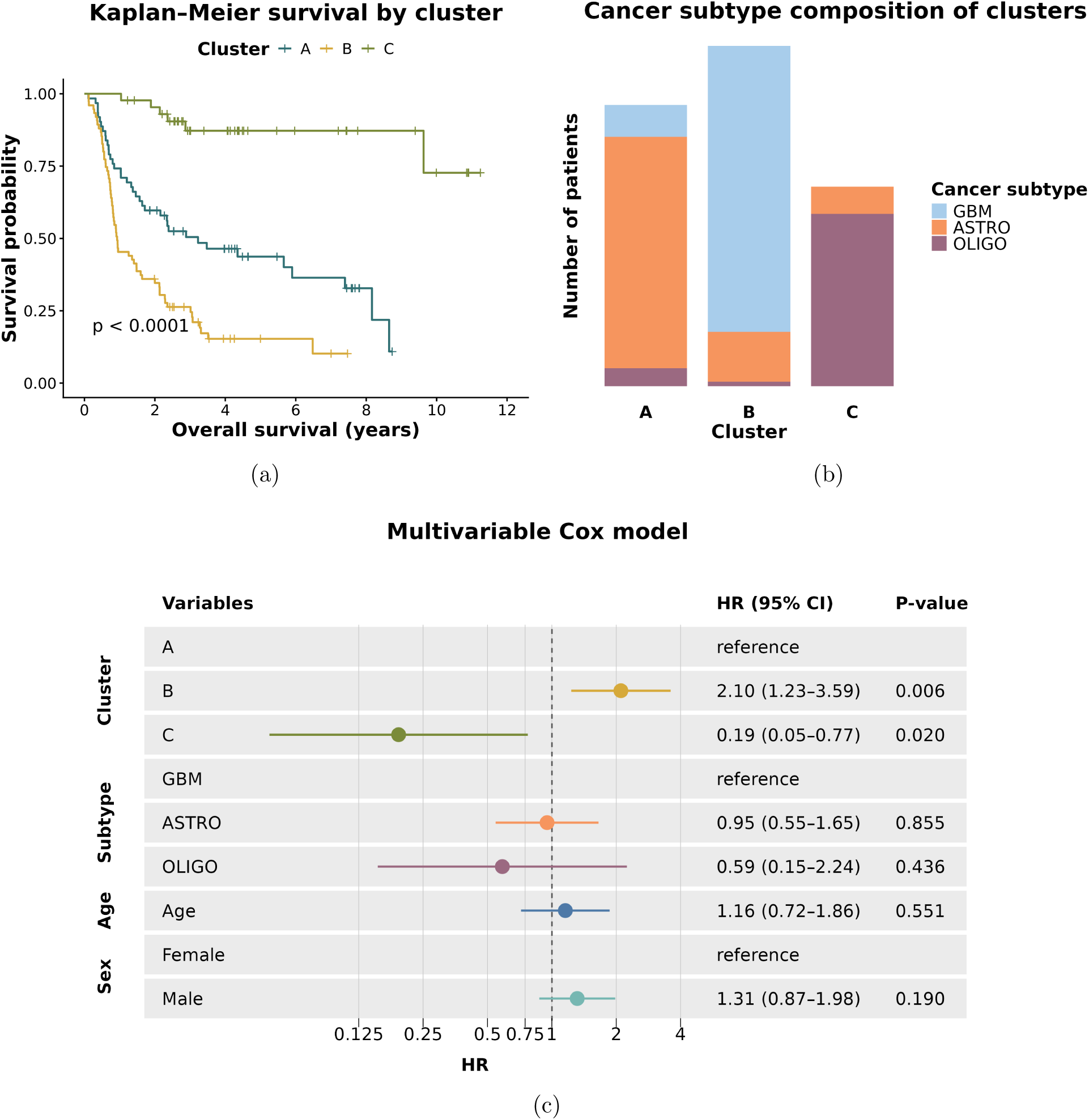
External survival stratification and WHO 2021 subtype composition of the TCGA-derived clusters in CGGA. a, Kaplan–Meier estimates of overall survival for the 181 CGGA patients assigned to Cluster A (*n* = 62), Cluster B (*n* = 75), or Cluster C (*n* = 44). Tick marks denote censoring; the displayed *p*-value is from the global log-rank test. **b,** Composition of each cluster by WHO 2021 tumour subtype: glioblastoma (GBM), astrocytoma (ASTRO), and oligodendroglioma (OLIGO). **c,** Hazard ratios (HRs) and 95% confidence intervals (CIs) from a multivariable Cox proportional-hazards model fitted in the CGGA validation cohort (*n* = 181; 106 events), including SNOWS cluster assignment, WHO 2021 tumour subtype, age, and sex (biological sex). Cluster A, glioblastoma, and female sex are the reference categories. Reported *p*-values are two-sided Wald-test values.

Although adjusted for WHO subtype, each cluster was dominated by one tumour type (Fig. 2b): Cluster A contained 51 astrocytomas, seven glioblastomas, and four oligodendrogliomas; Cluster B contained 63 glioblastomas, 11 astrocytomas, and one oligodendroglioma; and Cluster C contained 38 oligodendrogliomas and six astrocytomas. However, 29 of 181 patients (16.0%) were assigned outside the cluster dominated by their subtype, suggesting additional survival molecular heterogeneity beyond the WHO 2021 classification.

To assess incremental prognostic value, we compared a clinical multivariable Cox model including WHO subtype, age, and biological sex with a model additionally including SNOWS cluster assignment. Adding clusters increased the C-index from 0.691 to 0.712 (ΔC-index = 0.021; paired concordance test *p* = 3.4 *×* 10*^−^*^12^) and reduced IBS from 0.170 to 0.156 (ΔIBS = *−*0.014; Supplementary Table 2). The Cox log-likelihood also increased by 7.6 (likelihood-ratio test *p* = 4.9 *×* 10*^−^*^4^), supporting additional prognostic information beyond the clinical covariates. In the model including cluster assignment (Fig. 2c), Cluster B showed higher mortality risk than Cluster A (HR = 2.10, 95% CI 1.23–3.59, *p* = 0.006), whereas Cluster C showed lower risk (HR = 0.19, 95% CI 0.05–0.77, *p* = 0.020). In the same model, neither WHO subtype was statistically significant (ASTRO versus GBM: HR = 0.95, 95% CI 0.55–1.65, *p* = 0.855; OLIGO versus GBM: HR = 0.59, 95% CI 0.15–2.24, *p* = 0.436). These results suggest that SNOWS captures prognostic variation shared with WHO subtype while providing complementary survival information.

As a complementary analysis, we quantified the agreement between cluster assignment and clinical variables using normalised mutual information (NMI). Cluster assignment showed moderate, rather than complete, agreement with WHO subtype (NMI = 0.537), consistent with the subtype enrichment of each cluster but also with the presence of cross-subtype assignments. Agreement with age and sex was low (NMI = 0.089 and 0.016, respectively; Supplementary Table 2) indicating that the stratification was not primarily explained by these covariates. The combination of external survival separation, incremental performance across complementary metrics, and retained cluster effects after multivariable adjustment supports the interpretation of the SNOWS clusters as a prognostically informative refinement of established glioma classification.

### 3.2 Mutation profiles and network dependencies shape cluster assignments

Mutation-enrichment analysis in CGGA (Supplementary Table 4 and Fig. 7) revealed profiles consistent with the predominant WHO subtype in each cluster: *IDH1*, *TP53*, *ATRX* in astrocytoma-enriched Cluster A; *PTEN*, *NF1*, *EGFR* with IDH-wild-type status in glioblastoma-enriched Cluster B; and *IDH1*, *CIC*, *NOTCH1*, *FUBP1* in oligodendroglioma-enriched Cluster C [23, 24].

Each cluster is represented by a TCGA-derived BN (Fig. 3), allowing conditional dependencies among variables to contribute to patient assignment beyond marginal variable frequencies. To isolate this contribution, we compared BN-based assignments with an edge-free model retaining the fitted cluster priors and marginal variable distributions. Removing edges changed six of 181 CGGA assignments (3.3%). BN-based assignments yielded higher C-indices than the edge-free model, both for cluster-only Cox models (0.700 versus 0.688) and when including clinical covariates (0.712 versus 0.705; Supplementary Table 5). Thus, conditional dependencies refined a subset of assignments and were associated with improved prognostic discrimination.

**Figure 3:**
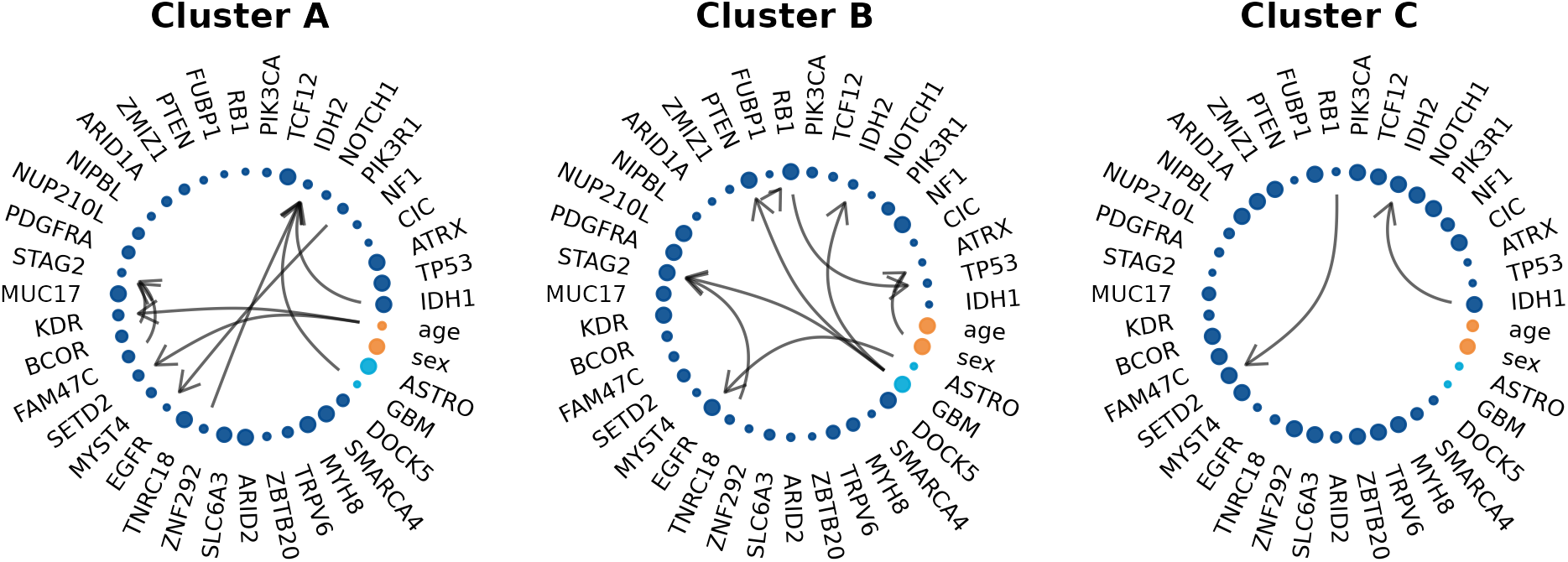
Cluster-specific Bayesian network structures learned in the TCGA glioma cohort. Nodes represent the molecular and clinical variables included in the model, and directed edges represent conditional dependencies. Node size is proportional to the frequency of the corresponding variable within the cluster, normalised across clusters for each variable. Dark blue nodes denote somatic mutations, light blue nodes denote WHO 2021 subtype, and orange nodes denote age and biological sex. Oligodendroglioma was used as the reference category for the subtype, therefore is not represented by a separate node.

For two GBM patients (CGGA_1343 and CGGA_P116), retaining BN dependencies changed assignment from the OLIGO-enriched Cluster C to the GBM-enriched Cluster B, restoring subtype concordance. The main contribution came from the *IDH2* node family, conditioned on the GBM indicator in Cluster B but on *IDH1* in Cluster C (+10.11 log-likelihood units; Supplementary Fig. 8). The remaining four cases were more informative because the dependencies supported cross-subtype assignments. CGGA_1415 and CGGA_J105 were both recorded as GBM and moved from Cluster C under the empty-BN classifier to the ASTRO-enriched Cluster A under the BN-based classifier. In both patients, the dominant contribution arose from the *IDH2* node family: in Cluster A, *IDH2* was modelled conditional on *IDH1*, *ZNF292*, and the GBM indicator, whereas in Cluster C it was conditioned only on *IDH1* (+10.11). Both patients were censored after 2719 and 3190 days of follow-up, respectively, which was compatible with the more favourable survival profile of Cluster A relative to the GBM-enriched Cluster B. CGGA_J35 was also recorded as GBM but moved from the Cluster B to Cluster A. The reassignment was driven primarily by the *ATRX* node family: *ATRX* was unconditioned in Cluster A but conditional on biological sex in Cluster B (+13.71). This patient remained censored at 2767 days, an outcome consistent with assignment away from the poor-survival Cluster B. CGGA_1985, recorded as ASTRO, moved from the ASTRO-enriched Cluster A to the OLIGO-enriched Cluster C. The change was driven predominantly by the *EGFR* node family: *EGFR* was unconditioned in Cluster C but conditional on *PIK3R1* in Cluster A (+9.69). Unlike the three subtype-discordant GBM cases, this patient experienced an event at 380 days despite assignment to the cluster with the most favourable OS profile. This case illustrates that, although BN-based cluster assignment captures cluster-level prognostic differences, it is not expected to predict the outcome of every individual patient with complete accuracy (Supplementary Fig. 9). Together, these cases show that the contribution of the BNs arises from differences in the context in which individual alterations are evaluated. For the four subtype-discordant patients, these conditional dependencies changed the relative support for clusters in ways that could not be explained by marginal mutation frequencies alone.

Cross-subtype assignment was not restricted to BN-sensitive patients. A further 25 patients remained assigned outside the cluster dominated by their WHO subtype after all edges were removed, indicating that their classification was driven primarily by marginal mutation patterns (Supplementary Fig. 10). These findings indicate that most subtype-discordant assignments were supported by mutation profiles resembling the assigned cluster rather than by dependency structures alone.

### 3.3 Comparative performance and generalisability of SNOWS

We next compared the incremental prognostic value of SNOWS with that of three alternative clustering approaches in the CGGA cohort (Supplementary Table 8): CANclust, the covariate-adjusted BN clustering framework without survival modelling [16]; a Bayesian network mixture model (BNMM), representing BN-based clustering without covariate adjustment [15] and using BiDAG for BN learning [25, 26]; and conventional *k*-means clustering. For each method, prognostic performance was evaluated by comparing a clinical-only Cox model (WHO 2021 tumour subtype, age, and biological sex) with a full model that additionally included cluster assignment. Among the evaluated methods, SNOWS achieved the highest full-model C-index of 0.712 (ΔC-index = 0.021, paired concordance test *p* = 3.41 *×* 10*^−^*^12^) and a log-likelihood gain of Δ*ℓ* = 7.6 (likelihood-ratio test *p* = 4.9 *×* 10*^−^*^4^). The corresponding full-model C-indices were 0.707 for CANclust (ΔC-index = 0.016, *p* = 1.70 *×* 10*^−^*^9^), 0.693 for BNMM (ΔC-index = 0.002, *p* = 0.0187), and 0.701 for k-means (ΔC-index = 0.010, *p* = 0.0469). Their respective log-likelihood gains were 6.0, 0.4 and 1.4 (*p* = 2.4 *×* 10*^−^*^3^, *p* = 0.67 and *p* = 0.24 based on likelihood-ratio tests). SNOWS also produced the lowest reported full-model IBS (0.156), closely followed by CANclust (0.160). The consistent difference between SNOWS and CANclust demonstrated a benefit from incorporating survival information during discovery. BNMM and k-means showed substantially stronger agreement with WHO subtype (NMI = 0.766 and 0.834, respectively) and smaller incremental survival associations.

To assess the generalisability of SNOWS, we evaluated its performance in TCGA pan-kidney and MD Anderson pan-myeloid cohorts using nested cross-validation (Supplementary Table 9). In pan-kidney cancer, adding SNOWS clusters increased the mean held-out C-index by 0.012 and reduced IBS by 0.0010, with C-index improvement in all five outer folds. In the pan-myeloid cohort, the gains were larger (ΔC-index = 0.079; ΔIBS = *−*0.022), again with C-index improvement in all five outer folds. These results show that SNOWS can provide additional prognostic information beyond glioma, although the magnitude of improvement varies across diseases.

## 4 Discussion

This study presents SNOWS, a probabilistic clustering framework in which cluster-specific BNs model dependencies among molecular and clinical covariates, while survival information guides the identification of prognostically distinct patient groups. In the principal application to adult-type diffuse glioma, SNOWS identified clusters that transferred from TCGA to an independent CGGA cohort and provided prognostic information complementary to WHO 2021 tumour subtype and the evaluated clinical covariates. The improvement in predictive performance was modest, but its consistency across analyses suggests that clinically established categories do not capture all survival information contained in the molecular profiles.

Despite adjustment for WHO subtype during discovery, SNOWS clusters remained strongly aligned with established glioma entities and their characteristic mutation profiles, reinforcing the biological foundation of WHO 2021 classification. However, this correspondence was incomplete, revealing patients with molecular configurations more characteristic of another prognostic group. SNOWS clusters should therefore be interpreted as complementary prognostic groups rather than new diagnostic entities.

Cluster-specific BNs capture conditional dependencies that may reveal molecular structure beyond individual alterations. The clearest example involves *IDH1* and *IDH2*, whose mutations are generally mutually exclusive in glioma, with the rarer *IDH2* mutations occurring more frequently in oligodendroglioma than astrocytoma [24, 27]. The involvement of IDH status and tumour subtype in the learned networks is therefore consistent with established glioma biology. In Cluster A, *ZNF292* also contributed to the network context of *IDH2*. Although *ZNF292* mutations have been reported at low frequency in lower-grade glioma [28], their specific relationship with *IDH2* remains unsupported and requires independent validation. A second informative dependency in Cluster A linked *EGFR* to *PIK3R1*. EGFR is a cell-surface receptor that can activate the PI3K signalling pathway, which promotes cell growth and survival, whereas *PIK3R1* encodes a regulatory component that helps control this pathway. Alterations in both genes have been reported in glioblastoma, and glioblastoma-associated *PIK3R1* mutations have been shown to increase PI3K signalling and promote tumour formation [29, 30].

In Cluster B, a dependency between the GBM indicator and *PTEN* was consistent with recurrent *PTEN* alterations in glioblastoma, this relationship is also biologically plausible because PTEN is a tumour suppressor that normally restrains PI3K signalling [23, 29]. Additional dependencies involved *TP53*, *RB1*, and *PTEN* ; these genes are associated with the p53, RB, and RTK–PI3K pathways, respectively, which are frequently disrupted in glioblastoma [29, 31]. Cluster B also showed a dependency between *ATRX* and bio-logical sex. Although a TCGA-based study reported sex-related differences in *ATRX* mutation clonality [32], it did not establish differences in mutation presence, so this association remains exploratory.

The sparse Cluster C network contained two displayed dependencies. One involved *IDH1* and *IDH2*, consistent with the established IDH-related molecular structure described above. The second dependency in Cluster C involved *SETD2* and *RB1*. *SETD2* mutations have been reported in molecularly defined oligodendroglioma and among its recurrent driver alterations, whereas *RB1* alterations are more characteristic of IDH-wild-type glioblastoma [23].

The empty-BN comparison showed that marginal mutation patterns explained most patient assignments, while conditional dependencies refined a smaller subset of patients with atypical molecular profiles. Cross-subtype assignments therefore arose from either resemblance to another cluster’s marginal mutation profile or additional network-specific information. These patterns reveal molecular heterogeneity beyond WHO subtypes without implying diagnostic misclassification.

The benchmarking analysis supports a similarly incremental interpretation of survival-informed clustering. SNOWS and CANclust performed relatively similarly, suggesting that incorporating survival during discovery refines rather than replaces the molecular structure identified by network-based clustering. By contrast, the stronger correspondence of BNMM and *k*-means with WHO subtype indicates that methods not explicitly adjusting for established clinical structure may preferentially recover the dominant diagnostic organisation already present in the data. Incorporating survival information while accounting for known clinical covariates may therefore help expose molecular variation associated with prognosis beyond this dominant signal. The differing prognostic gains in glioma, pan-kidney and pan-myeloid cohorts suggest that the benefits of survival-informed clustering depend on disease-specific molecular and clinical structure, rather than being uniform across malignancies.

Several limitations should be considered. The analyses were necessarily constrained by the molecular and clinical features available in each cohort. Differences in sequencing technologies, data-processing pipelines, clinical centres, and patient populations may introduce domain shifts that affect cluster assignment and prognostic performance. To reduce the risk of cohort-specific findings, we evaluated the TCGA-derived model in an independent CGGA cohort. More generally, incorporating additional molecular layers, treatment information, and richer clinical variables may reveal dependency structures not represented by the current analyses. Future extensions of SNOWS could incorporate transcriptomic, epigenomic, and proteomic features encoded as binary variables. Supporting continuous features would preserve more information and enable the integration of cellular abundances inferred from single-cell and bulk transcriptomic data, allowing molecular, clinical, and cellular dependencies to be modelled jointly.

Overall, SNOWS provides a general framework for identifying clinically relevant patient subgroups by integrating molecular dependency structure with survival information while accounting for established clinical covariates. Across the evaluated cohorts, this strategy captured prognostic variation not fully explained by the available clinical factors while retaining biologically coherent molecular organisation. In glioma, the analysis further showed that conditional dependencies can refine patient stratification beyond marginal mutation patterns alone. These findings support broader evaluation of survival-informed network clustering in larger, clinically harmonised cohorts with more comprehensive molecular and clinical characterisation.

## Data and code availability

SNOWS has been implemented as an R package. The package and all scripts required to reproduce the analyses and figures are available at https://github.com/sysbiomed/SNOWS. Analyses based on TCGA and CGGA can be reproduced using publicly available data. The pan-myeloid cohort is subject to data-access restrictions, and available as described in [16].

## Acknowledgements

This work was supported by Portuguese national funds through Fundação para a Cîencia e a Tecnologia, I.P. (FCT) under the PhD scholarship (2024.03867.BD) and projects UID/50021/2025, UID/PRR/50021/2025, UID/50022/2025, 2023.16283.ICDT, 2023.17447.ICDT. We also thank the all the Computational Biology Group at D-BSSE/ETHZ for hosting the visiting PhD research stay and providing a supportive research environment.

## Author contributions

Conceptualization: B.N.L., J.K., N.B., S.V.; Methodology: B.N.L., J.K., S.V.; Software: B.N.L.; Validation: B.N.L.; Formal analysis: B.N.L., S.V.; Investigation: B.N.L.; Resources: N.B., S.V.; Data curation: B.N.L., J.K.; Writing–original draft: B.N.L.; Writing–review & editing: B.N.L., J.K., N.B., S.V.; Visualization: B.N.L.; Supervision: J.K., N.B., S.V.; Project administration: J.K., N.B., S.V.; Funding acquisition: N.B., S.V.

## A Supplementary Material

### A.1 Study cohorts and analysis sets

The study comprised a TCGA glioma discovery cohort, an independent CGGA glioma validation cohort, a TCGA pan-kidney cohort, and a pan-myeloid malignancy cohort from MD Anderson Cancer Center. The glioma cohorts included glioblastoma (GBM), astrocytoma (ASTRO), and oligodendroglioma (OLIGO). TCGA-KIPAN comprised kidney chromophobe carcinoma (KICH), kidney renal papillary cell carcinoma (KIRP), and kidney renal clear cell carcinoma (KIRC). The pan-myeloid cohort included acute myeloid leukaemia (AML), chronic myelomonocytic leukaemia (CMML), myelodysplastic syndromes (MDS), and myeloproliferative neoplasms (MPN). Cohort sizes and clinical characteristics are detailed in Supplementary Table 1. TCGA glioma was used to select the molecular feature set, tune the glioma model, and estimate the final cluster-specific networks. CGGA was used for external assignment and subsequent evaluation of the prognostic and molecular characteristics of the transferred clusters. The pan-kidney and pan-myeloid cohorts were evaluated using nested cross-validation.

**Table 1:** Clinical and molecular characteristics of the study cohorts. Values are median (interquartile range) or number (percentage). OS, overall survival; AF, altered features per patient. OS is reported in days for TCGA glioma, CGGA glioma, and TCGA-KIPAN, and in months for the MDA pan-myeloid cohort. AF denotes mutated genes for the glioma and pan-kidney cohorts and mutated genes plus cytogenetic profiles for the pan-myeloid cohort.

| Characteristic | TCGA glioma | CGGA glioma | TCGA-KIPAN | MDA pan-myeloid |
|---|---|---|---|---|
| Patients, *n* | 950 | 181 | 898 | 1320 |
| Female, *n* (%) | 397 (41.8%) | 75 (41.4%) | 307 (34.2%) | 531 (40.2%) |
| Male, *n* (%) | 553 (58.2%) | 106 (58.6%) | 591 (65.8%) | 789 (59.8%) |
| Subtypes, *n* (%) | GBM: 502 (52.8%)<br>ASTRO: 281 (29.6%)<br>OLIGO: 167 (17.6%) | GBM: 70 (38.7%)<br>ASTRO: 68 (37.6%)<br>OLIGO: 43 (23.8%) | KICH: 112 (12.5%)<br>KIRP: 254 (28.3%)<br>KIRC: 532 (59.2%) | AML: 543 (41.1%)<br>CMML: 118 (8.9%)<br>MDS: 490 (37.1%)<br>MPN: 169 (12.8%) |
| Events, *n* (%) | 496 (52.2%) | 106 (58.6%) | 226 (25.2%) | 725 (54.9%) |
| Censored, *n* (%) | 454 (47.8%) | 75 (41.4%) | 672 (74.8%) | 595 (45.1%) |
| OS, median (IQR) | 511.5 (233.2–919.0) days | 837.0 (296.0–1512.0) days | 1073.0 (506.2–1896.0) days | 22.5 (8.8–48.5) months |
| AF, median (IQR) | 2.0 (0.0–3.0) | 2.0 (1.0–3.0) | 1.0 (0.0–2.0) | 3.0 (2.0–4.0) |

### A.2 Evaluation metrics

#### A.2.1 Likelihood-ratio analysis

The prognostic information provided by cluster assignment was evaluated using likelihood-ratio analysis within Cox proportional hazards models. For each clustering approach, a clinical Cox model was compared with an extended model that additionally included cluster assignment as a categorical variable.

Let *ℓ*_clinical_ and *ℓ*_full_ denote the maximised Cox partial log-likelihoods of the clinical and full models, respectively. The improvement in model fit attributable to cluster assignment was quantified as the log-likelihood gain

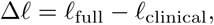

and statistical significance was assessed using the corresponding likelihood-ratio test (the test statistic is 2Δ*ℓ*, which is compared to the asymptotic *χ*^2^ null distribution with *K −* 1 degrees of freedom for *K* clusters).

#### A.2.2 C-index

Harrell’s concordance index (C-index) [33] was used to evaluate the predictive accuracy of survival models by assessing the model’s ability to predict higher relative risks to individuals whose event occurs first. It checks whether the model can correctly predict who has a higher risk for all possible pairs of individuals. A C-index of 1 means perfect concordance between the predicted and observed survival outcomes, while 0.5 indicates random guessing. In a survival analysis, certain pairs of observations can not be directly compared. For a pair to be comparable, either both individuals must have experienced the event (death), or one individual must have experienced the event while the other was lost to follow-up after the event occurred in the first. As a result, not all patient pairs can be evaluated, and the C-index is calculated only over comparable pairs. The survival package (version 3.8-3) in R was used to calculate the C-index.

To assess whether adding cluster assignment improved concordance beyond the clinical-only model, the C-indices of the clinical and full models were additionally compared using a paired concordance test. The comparison was performed using the head-to-head U-statistic implemented in Hmisc::rcorrp.cens (Hmisc version 5.2-5), which evaluates the two model predictions on the same comparable patient pairs. Statistical significance was assessed using the corresponding two-sided normal test.

#### A.2.3 Integrated Brier Score (IBS)

The Brier Score (BS) [34] quantifies the accuracy of probabilistic predictions by measuring the average difference between observed outcomes and predicted probabilities. Without censored data, it is calculated as the squared difference between a patient’s observed status and the predicted probability. To handle censoring, the survival Brier score relies on inverse probability of censoring weights (IPCW), following the approach proposed by Graf et al. [35]. These weights adjust each patient’s contribution according to the estimated censoring survival function, allowing prediction accuracy to be evaluated while accounting for right-censored observations. To evaluate the BS over all time points, the Integrated Brier Score (IBS) is used, which averages the BS over a range of times. This integral form ensures that the evaluation of the BS is not restricted to a specific time point, providing a more comprehensive measure of prediction accuracy. A lower IBS value indicates better model performance, reflecting more accurate predictions of survival probabilities across the considered time range. The pec package (version 2025.06.24) in R was used to implement these calculations.

#### A.2.4 Normalised mutual information (NMI)

The normalised mutual information (NMI) was used to evaluate the pairwise relationship between cluster assignments and individual clinical variables. NMI derives from the concept of mutual information introduced in information theory by Shannon [36] and has been adopted as an external measure of agreement between cluster partitions [37]. In this context, it quantifies the extent to which knowledge of a clinical variable reduces uncertainty about a patient’s cluster assignment. NMI values close to 0 indicate little dependence between the variables, whereas higher values indicate stronger agreement or overlap. NMI was calculated using the aricode package (version 1.0.3) in R.

#### A.2.5 Pseudo-*R*^2^

As a complementary analysis, McFadden’s pseudo-*R*^2^ was used to assess whether cluster assignments could be explained by clinical variables considered jointly [38]. Cluster membership was modelled as the outcome in a multinomial logistic regression, with clinical variables used as predictors. McFadden’s pseudo-*R*^2^ compares the likelihood of this model with that of a baseline model containing no predictors, thereby quantifying the extent to which clinical variables allow cluster assignments to be predicted. Lower values indicate that cluster assignments are less predictable from the evaluated clinical variables, suggesting greater independence from known clinical factors.

All analyses were performed in R version 4.5.1.

### A.3 Model selection and hyperparameter sensitivity

#### A.3.1 Selection of the number of clusters

To select the number of clusters, we fitted unsupervised Bayesian network mixture models (equivalent to Bernoulli mixture models) across candidate cluster numbers and pseudocount values. For each pseudocount, the preferred number of clusters was identified using the Bayesian information criterion (BIC), with lower values indicating better fit. Fig. 4 show the differences in BIC relative to the minimum obtained at each pseudocount. The analysis consistently supported three clusters for TCGA glioma and TCGA-KIPAN and six clusters for MDA pan-myeloid across the evaluated pseudocount values.

**Figure 4:**
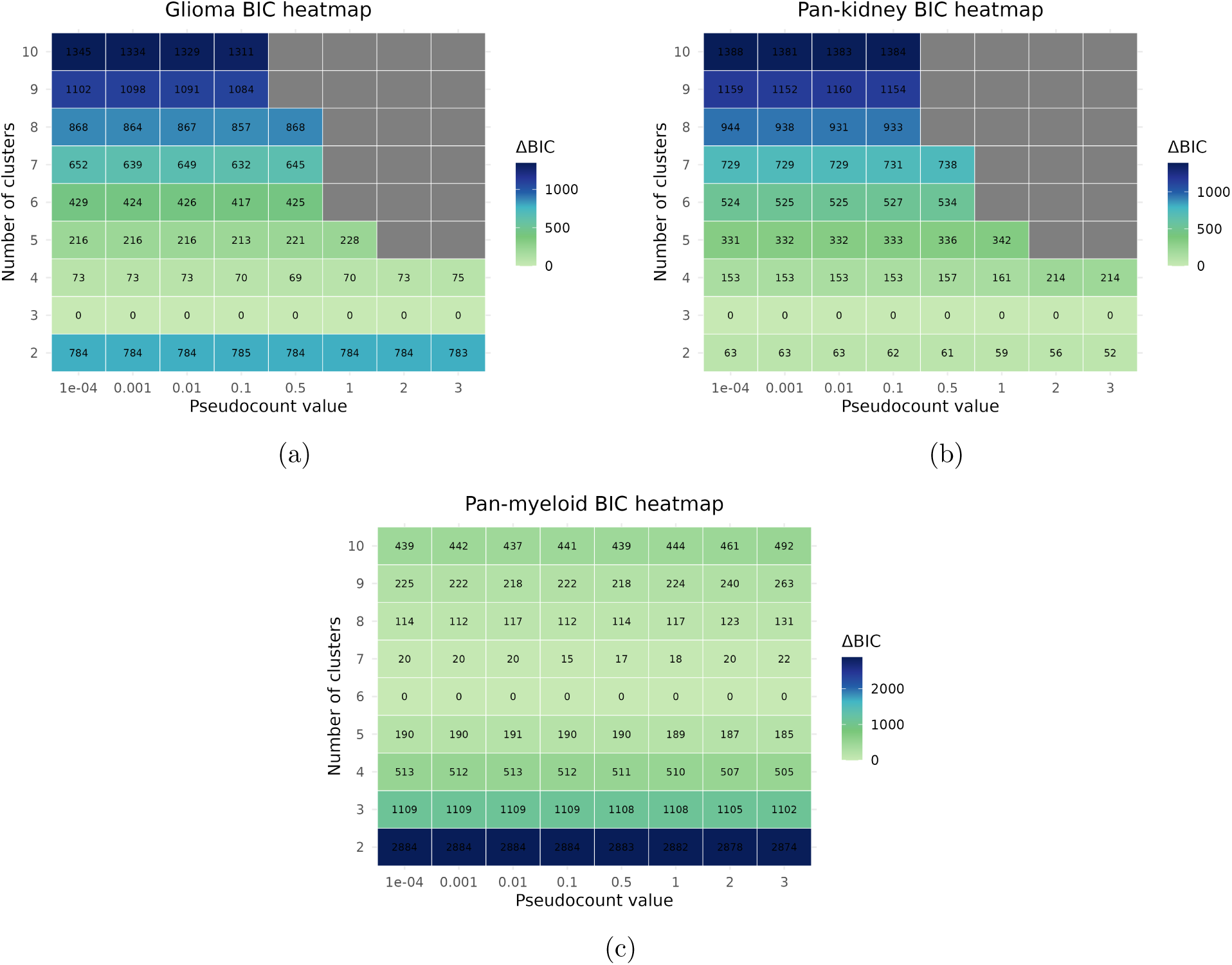
Selection of the number of clusters using BIC. ΔBIC was calculated relative to the minimum BIC across the evaluated numbers of clusters for each pseudocount value, with lower values indicating better fit. **a,** TCGA glioma (*K* = 3). **b,** TCGA-KIPAN (*K* = 3). **c,** MDA pan-myeloid (*K* = 6). The selected number of clusters was consistent across the evaluated pseudocount values.

#### A.3.2 Sensitivity to the BN pseudocount and survival weight

We evaluated the sensitivity of the glioma model to the Bayesian network pseudocount, *χ*, and the survival weight, *α*, using five-fold cross-validation in the TCGA discovery cohort. The primary selection criterion was the mean ΔC-index, calculated as the difference between the C-index of the clinical-plus-cluster model and that of the clinical-only model. This measures the additional prognostic discrimination provided by the SNOWS-derived clusters. Mean ΔIBS and the cluster-only C-index were examined as complementary measures of prognostic performance, while the mean minimum cluster size was used to identify parameter settings for which one of the three clusters became very small or collapsed.

The results supported the use of a small, non-zero pseudocount (Supplementary Fig. 5). Although *χ* = 0 generally preserved large clusters, it produced less favourable incremental prognostic performance. Increasing the pseudocount did not consistently improve the minimum cluster size and was generally associated with weaker ΔC-index, ΔIBS, and cluster-only C-index values. In contrast, *χ* = 0.001 retained a region at low values of *α* in which all three clusters remained represented while the resulting partition showed favourable prognostic performance. This low pseudocount places limited weight on prior smoothing of the Bayesian network parameters, allowing the molecular structure of the data to remain influential during cluster discovery.

**Figure 5:**
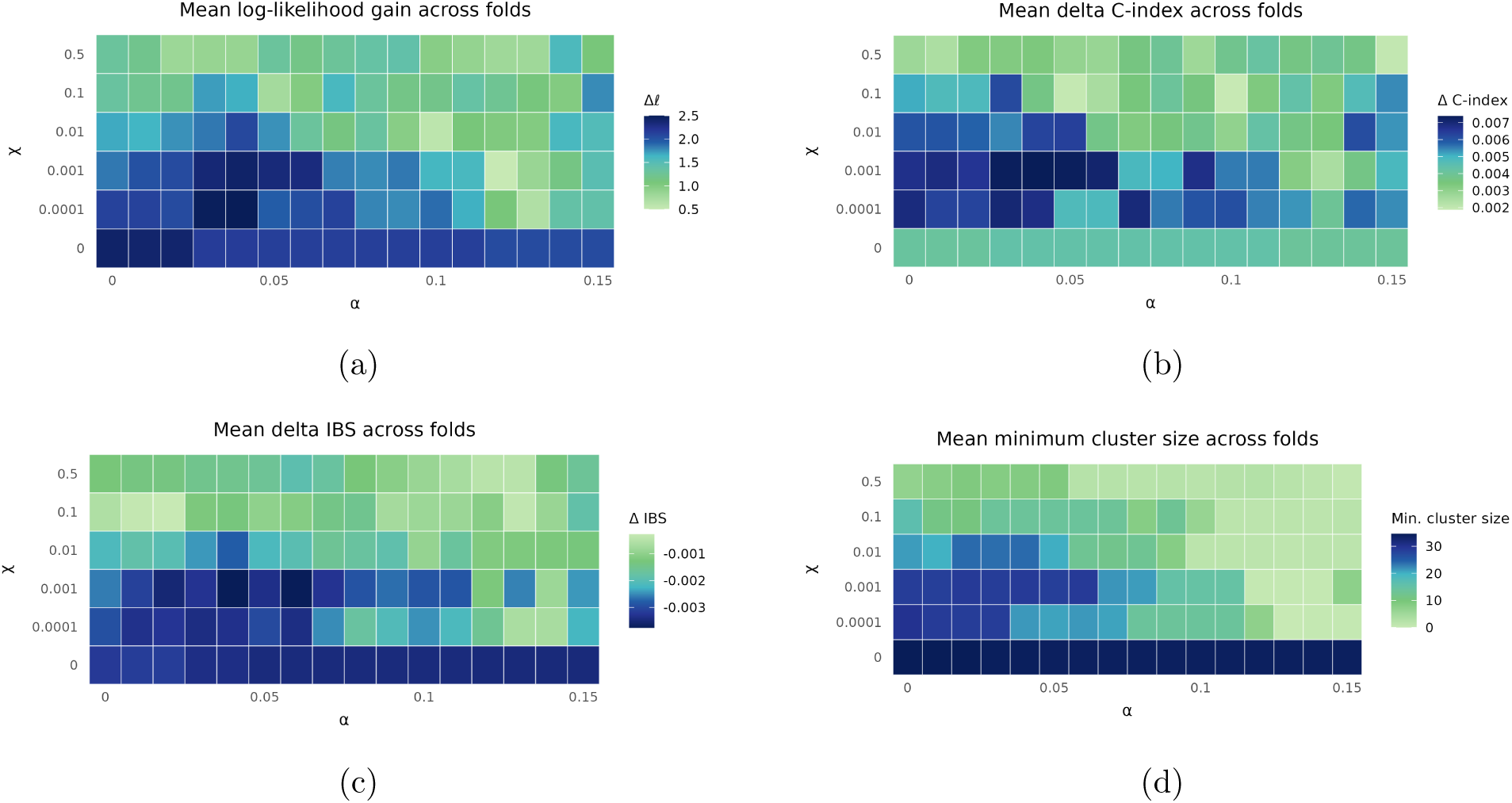
Sensitivity of the glioma model to the Bayesian network pseudocount and survival weight. Heatmaps show results averaged across five cross-validation folds in the TCGA glioma discovery cohort for each combination of the pseudocount *χ* and survival weight *α*. **a,** The log-likelihood gain (Δ*ℓ*) quantifies the improvement in Cox-model fit obtained after adding cluster assignment to the clinical model. **b,** ΔC-index was calculated as the C-index of the clinical-plus-cluster model minus that of the clinical-only model, with larger positive values indicating greater incremental prognostic discrimination. **c,** ΔIBS was calculated analogously, with more negative values indicating lower prediction error. **d,** The minimum cluster size indicates whether one of the three clusters became very small or collapsed. The selected setting, *χ* = 0.001 and *α* = 0.04, combined favourable model fit and prognostic performance with preservation of all three clusters.

For *χ* = 0.001, increasing *α* from zero improved the mean ΔC-index, suggesting that a small contribution from survival information enhanced the prognostic relevance of the learned clusters. We selected *α* = 0.04 because it lay within the region of highest mean ΔC-index while also showing a favourable mean ΔIBS and cluster-only C-index. Importantly, the minimum cluster size remained comparatively large at this setting. Further increases in *α* did not provide a consistent improvement in prognostic performance and progressively reduced the size of the smallest cluster. The selected combination, *χ* = 0.001 and *α* = 0.04, therefore balanced the additional prognostic information introduced by survival-guided discovery with preservation of a meaningful three-cluster solution.

### A.4 External prognostic validation in CGGA

#### A.4.1 Survival according to WHO 2021 subtype

WHO 2021 tumour subtype was strongly associated with overall survival in CGGA (Supplementary Fig. 6; global log-rank *p <* 0.0001). Glioblastoma had the poorest observed survival, oligodendroglioma the most favourable survival, and astrocytoma an intermediate profile. This expected hierarchy provides context for interpreting the close, but incomplete, correspondence between WHO subtype and the transferred clusters.

**Figure 6:**
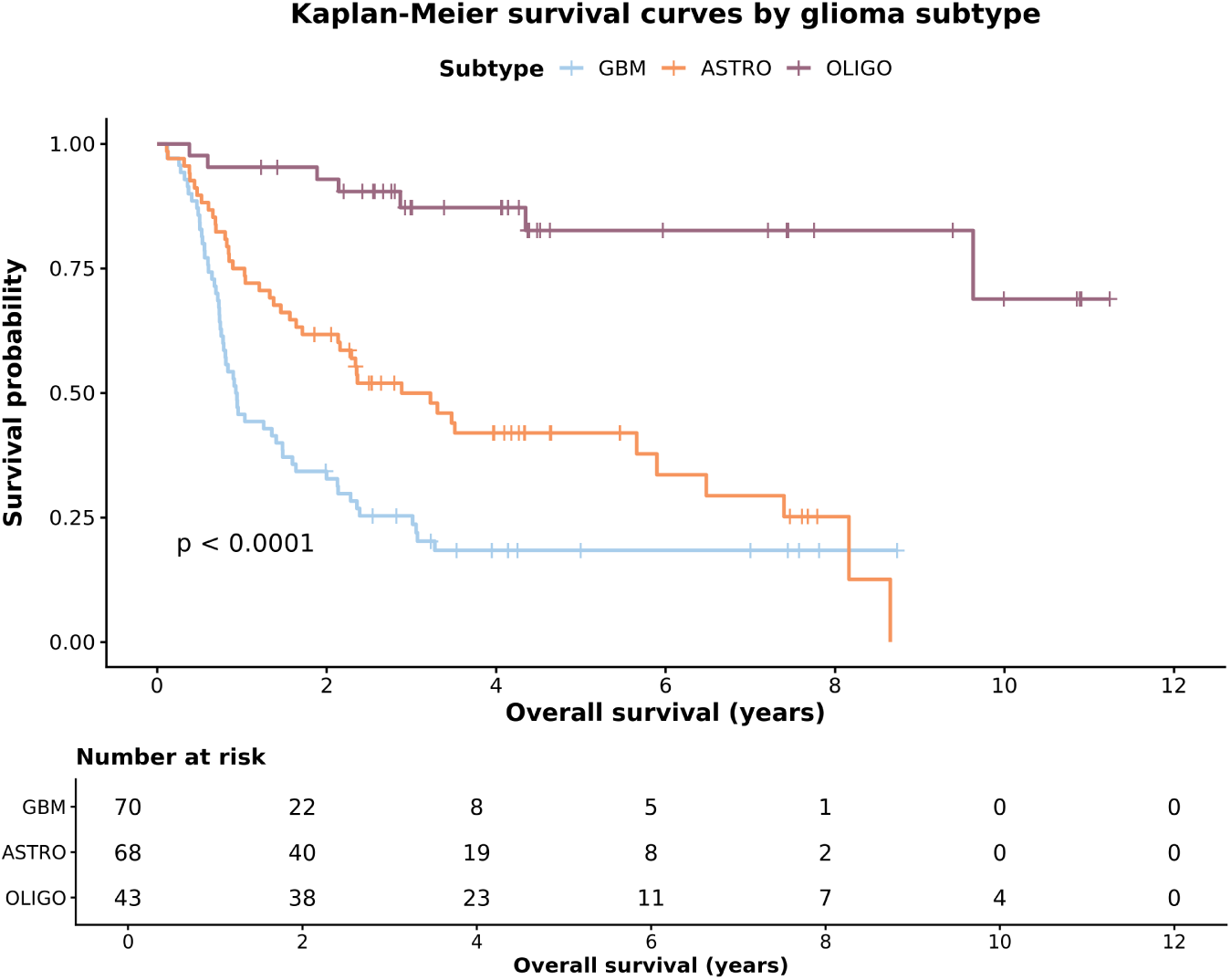
Overall survival according to WHO 2021 glioma subtype in CGGA. Kaplan–Meier estimates are shown for glioblastoma (*n* = 70), astrocytoma (*n* = 68), and oligodendroglioma (*n* = 43). Tick marks denote censoring; the displayed *p*-value is from the global log-rank test.

#### A.4.2 Prognostic performance of SNOWS clusters

Supplementary Table 2 reports the descriptive performance and cluster–covariate association measures used in the primary CGGA analysis. The cluster-only and subtype-only models contain only the indicated categorical variable. The clinical-only model contains the prespecified clinical covariates, and the full model adds cluster assignment to those covariates.

#### A.4.3 Sensitivity to clinical–molecular IDH discordance

When analysing mutation frequencies across WHO subtype–cluster combinations, we found that two patients classified as oligodendroglioma and 19 classified as astrocytoma had no qualifying *IDH1* or *IDH2* alteration in the processed mutation data, despite both subtypes being defined as IDH-mutant under the WHO 2021 classification. This created a discrepancy between the clinical annotation, which recorded these patients as IDH-mutant, and the independently processed mutation data. The available data do not establish which representation more accurately reflects the underlying tumour.

**Table 2:** Prognostic metrics and association with clinical variables for the transferred SNOWS clusters in CGGA. ΔC-index and ΔIBS compare the full model with the clinical-only model.

| Metric | Value |
|---|---|
| Log-likelihood gain (Δℓ) | 7.6 |
| Likelihood-ratio test *p*-value | 4.9 × 10<sup>−4</sup> |
| C-index, clinical-only model | 0.691 |
| C-index, cluster-only model | 0.700 |
| C-index, subtype-only model | 0.690 |
| C-index, full model | 0.712 |
| ΔC-index, full versus clinical-only | 0.021 |
| Paired concordance test *p*-value | 3.4 × 10<sup>−12</sup> |
| IBS, clinical-only model | 0.170 |
| IBS, cluster-only model | 0.157 |
| IBS, subtype-only model | 0.171 |
| IBS, full model | 0.156 |
| ΔIBS, full versus clinical-only | -0.014 |
| McFadden pseudo-*R*<sup>2</sup> | 0.561 |
| Cluster novelty index | 0.439 |
| NMI with WHO 2021 subtype | 0.537 |
| NMI with sex | 0.016 |
| NMI with age | 0.089 |
| Cluster size range | 44–75 |
| Number of patients | 181 |

SNOWS does not require either representation to be selected as correct; instead, both the subtype and the observed mutation profile contribute to the evaluation of cluster membership, allowing patients to be assigned according to the joint support of their complete clinical and mutational profile. Some of these patients were assigned to the GBM-enriched Cluster B, demonstrating that assignments were not determined solely by recorded subtype and could instead evaluate the wider mutational and clinical configuration. Excluding these 21 patients did not remove the incremental prognostic improvement (ΔC-index = 0.027 in the remaining 160 patients versus 0.021 in the full cohort; Supplementary Table 3). Although these estimates are not directly comparable because cohort composition and comparable survival pairs differ, analysis suggests that the observed improvement was not predominantly driven by these discordant cases.

### A.5 Molecular characterisation and BN contributions

#### A.5.1 Mutation-enrichment analysis

For each of the 35 TCGA-selected genes, mutation frequency within a CGGA cluster was compared with the frequency among all patients assigned to the other two clusters using a two-sided Fisher’s exact test. Benjamini–Hochberg correction was applied separately within each cluster across the 35 genes. Supplementary Table 4 reports genes with adjusted *p <* 0.05.

#### A.5.2 BN-based versus edge-free assignments

The empty-BN classifier retained the fitted cluster priors and cluster-specific marginal distributions but removed every learned edge. This comparison isolates the contribution of conditional dependency terms to external assignment. BN-based and empty-BN performance is summarised in Supplementary Table 5.

**Table 3:** Sensitivity analysis after exclusion of patients with clinical–molecular IDH discordance. Patients assigned to an IDH-mutant WHO 2021 tumour type without a qualifying *IDH1* or *IDH2* alteration in the processed mutation data were excluded. The clinical model included WHO 2021 tumour subtype, age, and biological sex, and the full model additionally included SNOWS cluster assignment.

| Metric | Value |
|---|---|
| Log-likelihood gain (Δℓ) | 8.1 |
| Likelihood-ratio test *p*-value | 3.2 × 10<sup>−4</sup> |
| C-index, clinical-only model | 0.699 |
| C-index, cluster-only model | 0.715 |
| C-index, full model | 0.726 |
| ΔC-index, full versus clinical-only | 0.027 |
| IBS, clinical-only model | 0.166 |
| IBS, cluster-only model | 0.150 |
| IBS, full model | 0.144 |
| ΔIBS, full versus clinical-only | -0.022 |
| Cluster size range | 42–63 |
| Number of patients | 160 |

**Table 4:** Genes significantly enriched or depleted in the transferred CGGA clusters. Frequencies are the proportions of patients with a qualifying non-silent mutation inside and outside the indicated cluster. OR, odds ratio. Adjusted *p*-values are from two-sided Fisher’s exact tests followed by Benjamini–Hochberg correction within each cluster.

| Cluster | Gene | Direction | Frequency in cluster | Frequency outside cluster | OR | Adjusted *p*-value |
|---|---|---|---|---|---|---|
| A | *IDH1* | Enriched | 77.4% | 32.8% | 6.95 | 1.2 × 10<sup>−7</sup> |
|  | *TP53* | Enriched | 95.2% | 28.6% | 48.00 | 3.4 × 10<sup>−18</sup> |
|  | *ATRX* | Enriched | 80.6% | 5.0% | 74.70 | 9.8 × 10<sup>−25</sup> |
|  | *CIC* | Depleted | 0.0% | 20.2% | 0.00 | 2.8 × 10<sup>−4</sup> |
| B | *IDH1* | Depleted | 0.0% | 82.1% | 0.00 | 1.6 × 10<sup>−31</sup> |
|  | *ATRX* | Depleted | 6.7% | 48.1% | 0.08 | 8.8 × 10<sup>−9</sup> |
|  | *CIC* | Depleted | 1.3% | 21.7% | 0.05 | 3.1 × 10<sup>−4</sup> |
|  | *NF1* | Enriched | 14.7% | 1.9% | 8.84 | 1.2 × 10<sup>−2</sup> |
|  | *NOTCH1* | Depleted | 0.0% | 15.1% | 0.00 | 1.5 × 10<sup>−3</sup> |
|  | *PTEN* | Enriched | 18.7% | 1.9% | 11.79 | 9.2 × 10<sup>−4</sup> |
|  | *EGFR* | Enriched | 10.7% | 0.9% | 12.39 | 2.0 × 10<sup>−2</sup> |
| C | *IDH1* | Enriched | 88.6% | 35.0% | 14.25 | 2.5 × 10<sup>−9</sup> |
|  | *TP53* | Depleted | 4.5% | 66.4% | 0.02 | 1.0 × 10<sup>−12</sup> |
|  | *ATRX* | Depleted | 2.3% | 40.1% | 0.04 | 1.5 × 10<sup>−6</sup> |
|  | *CIC* | Enriched | 52.3% | 0.7% | 142.78 | 1.9 × 10<sup>−14</sup> |
|  | *NOTCH1* | Enriched | 25.0% | 3.6% | 8.66 | 7.0 × 10<sup>−4</sup> |
|  | *FUBP1* | Enriched | 11.4% | 0.7% | 17.10 | 2.0 × 10<sup>−2</sup> |

**Table 5:** Descriptive comparison of BN-based and empty-BN assignments in CGGA. The empty-BN classifier removes all learned edges while retaining fitted cluster priors and cluster-specific marginal mutation probabilities. ΔC-index compares the full clinical model with the clinical-only model.

| Metric | BN-based | Empty-BN |
|---|---|---|
| Log-likelihood gain (Δℓ) | 7.6 | 7.1 |
| Likelihood-ratio test *p*-value | 4.9 × 10<sup>−4</sup> | 8.0 × 10<sup>−4</sup> |
| Cluster-only C-index | 0.700 | 0.688 |
| Full-model C-index | 0.712 | 0.705 |
| ΔC-index, full versus clinical-only | 0.021 | 0.014 |

**Figure 7:**
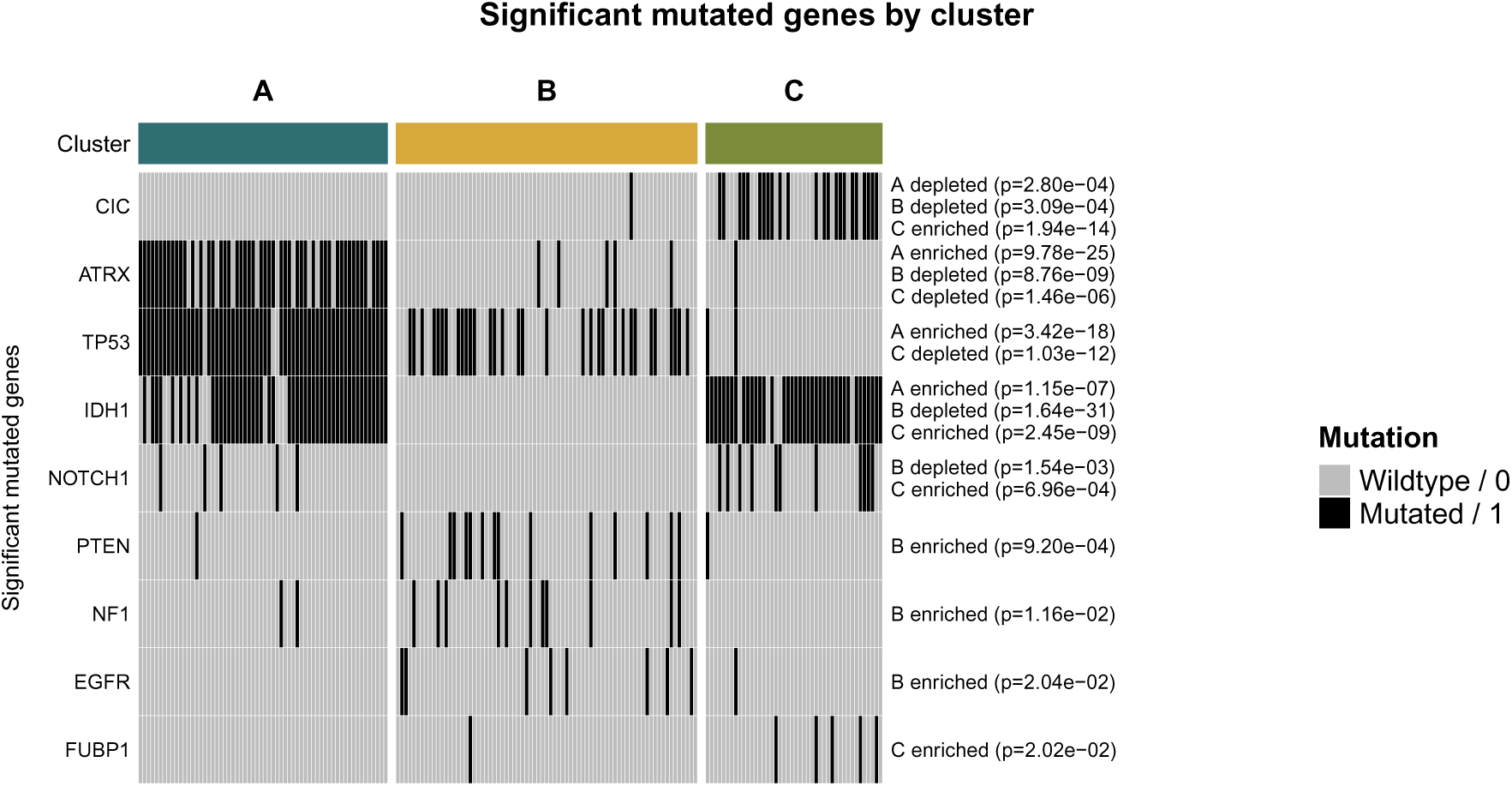
Cluster-associated mutation patterns in the CGGA validation cohort. Columns represent patients ordered by assigned cluster and rows represent genes that were significantly enriched or depleted in at least one cluster using two-sided Fisher’s exact tests with Benjamini–Hochberg correction. Black indicates a qualifying non-silent mutation and grey indicates no qualifying mutation. Text at right reports the direction and adjusted *p*-value of significant comparisons. Exact mutation frequencies, odds ratios, raw *p*-values, and adjusted *p*-values are provided in Supplementary Table 4.

#### A.5.3 BN-sensitive and subtype concordance

A patient was defined as BN-sensitive if removing all BN edges changed their assigned cluster. Subtype-discordant assignment refers to assignment to a cluster whose predominant WHO subtype differs from the patient’s recorded subtype. Six patients were BN-sensitive, of whom two were subtype-concordant and four subtype-discordant (Supplementary Table 6). Among the 175 patients whose assignments were unchanged, 25 were nevertheless assigned outside the cluster dominated by their recorded subtype. This distinction identifies cross-subtype assignments that persisted without network edges and those specifically altered by conditional dependencies.

**Table 6:** BN sensitivity and subtype concordance in CGGA. A patient was defined as BN-sensitive if removing all BN edges changed their assigned cluster. Subtype-discordant assignment refers to assignment to a cluster whose predominant WHO subtype differs from the patient’s recorded subtype.

|  | BN-sensitive | Not BN-sensitive | Total |
|---|---|---|---|
| Subtype-concordant | 2 (1.10%) | 150 (82.87%) | 152 (83.98%) |
| Subtype-discordant | 4 (2.21%) | 25 (13.81%) | 29 (16.02%) |
| Total | 6 (3.31%) | 175 (96.69%) | 181 (100.00%) |

#### A.5.4 Local node-family score decomposition

For each BN-sensitive patient, we compared the BN-based score of the selected cluster with the BN-based score of the cluster selected by the empty-BN classifier. The difference was decomposed into local node-family terms, where a node family comprises a child node and its fitted parent set. Positive values favour the BN-based–selected cluster.

**Table 7:** Clinical metadata and assignments of the six BN-sensitive CGGA patients. Patients are grouped by whether their BN-based assignment was concordant with the predominant WHO subtype of the assigned cluster. Follow-up is reported in days.

| Group | Sample | WHO subtype | Empty-BN | BN-based | Predominant subtype | Follow-up | Status |
|---|---|---|---|---|---|---|---|
| Subtype-concordant | CGGA_1343 | GBM | C | B | GBM | 94 | Event |
|  | CGGA_P116 | GBM | C | B | GBM | 305 | Event |
| Subtype-discordant | CGGA_1415 | GBM | C | A | ASTRO | 2719 | Censored |
|  | CGGA_1985 | ASTRO | A | C | OLIGO | 380 | Event |
|  | CGGA_J105 | GBM | C | A | ASTRO | 3190 | Censored |
|  | CGGA_J35 | GBM | B | A | ASTRO | 2767 | Censored |

For CGGA_1343 and CGGA_P116, the largest positive term was the *IDH2* family (+10.11 in each patient), followed by smaller contributions from *PTEN*, *IDH1*, *TP53*, *RB1*, *PDGFRA*, *CIC*, or *FUBP1* (Supplementary Fig. 8). For the subtype-discordant patients, the largest terms were the *IDH2* family for CGGA_1415 and CGGA_J105 (+10.11), the *EGFR* family for CGGA_1985 (+9.69), and the *ATRX* family for CGGA_J35 (+13.71), with a secondary *TP53* term for CGGA_J35 (+2.15; Supplementary Fig. 9).

**Figure 8:**
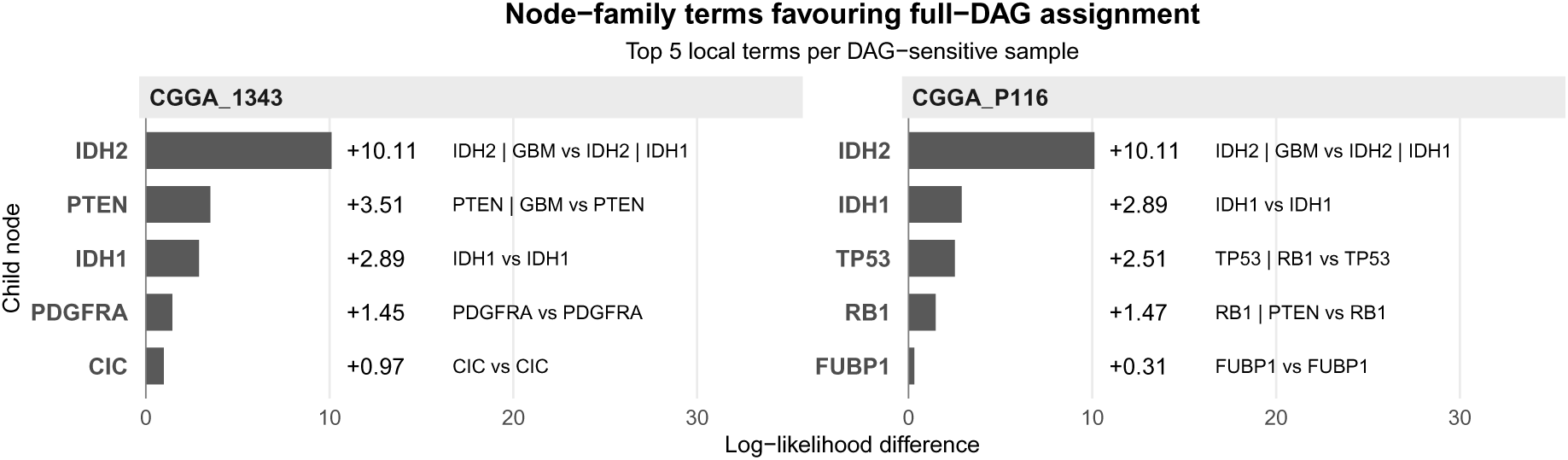
Largest positive node-family terms for BN-sensitive, subtype-concordant CGGA patients. Bars show the local log-likelihood difference between the BN-based–selected cluster and the empty-BN-selected cluster after both candidate clusters are evaluated with their BN-based classifiers. Only the five largest positive terms per patient are displayed. Labels identify the child and parent set in the two compared clusters.

#### A.5.5 Mutation profiles across WHO subtype–cluster combinations

Among the 29 subtype-discordant patients, 25 were not BN-sensitive, indicating that their cross-subtype assignments did not require the learned edges. Supplementary Fig. 10 summarises frequencies of the nine genes that were significantly enriched or depleted at cluster level.

### A.6 Benchmarking and generalisability

#### A.6.1 Benchmarking in CGGA

The prognostic performance of SNOWS was compared with three alternative clustering approaches (CAN-clust, BNMM, and k-means) in the CGGA validation cohort (Supplementary Table 8). For each method, Cox proportional hazards models were fitted using either the clinical covariates alone (WHO 2021 tumour subtype, age, and biological sex) or the same covariates together with cluster assignment (full model).

**Figure 9:**
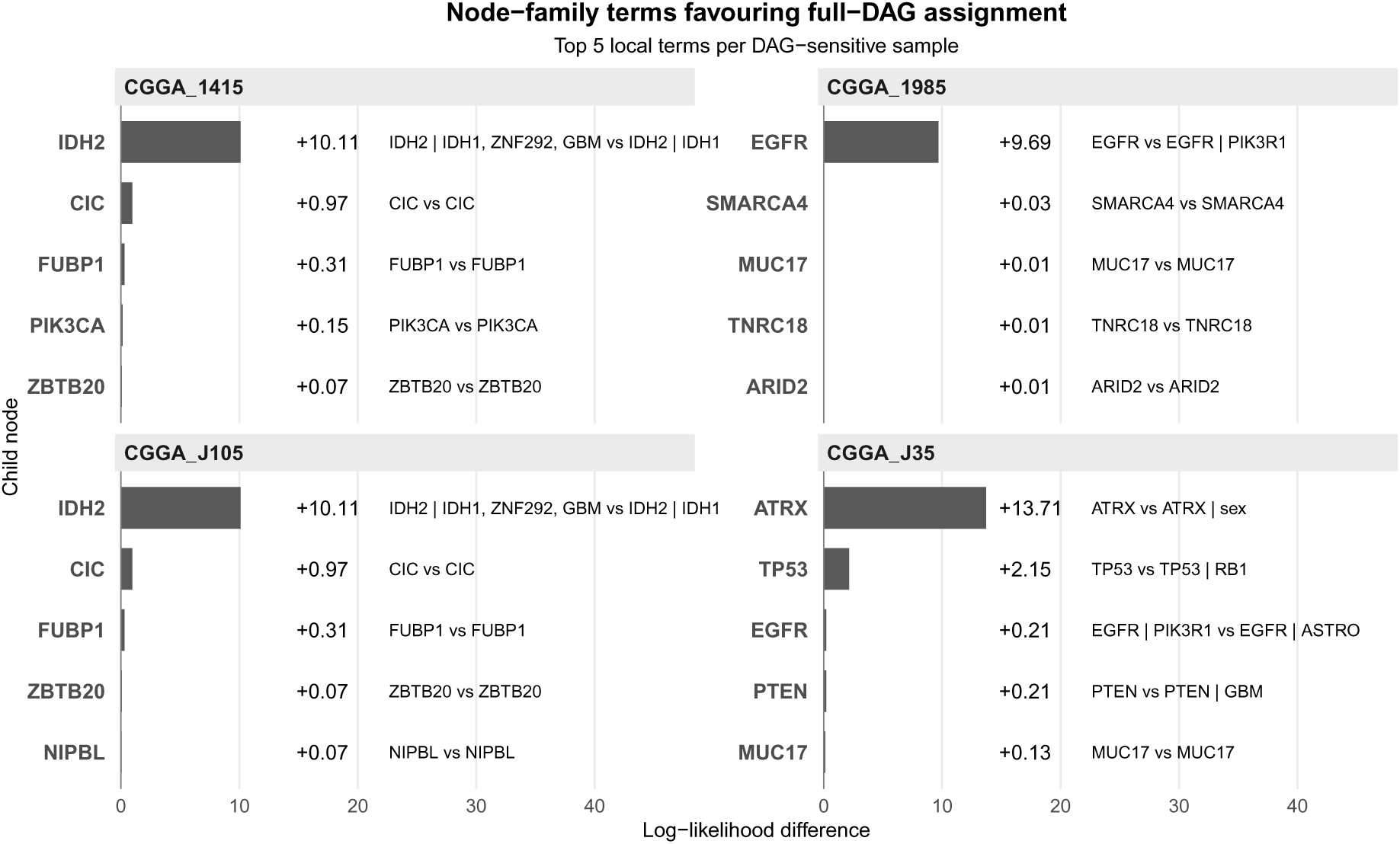
Largest positive node-family terms for BN-sensitive, subtype-discordant CGGA patients. Bars show the local log-likelihood difference between the BN-based–selected cluster and the empty-BN–selected cluster after both candidate clusters are evaluated with their BN-based classifiers. Only the five largest positive terms per patient are displayed.

**Figure 10:**
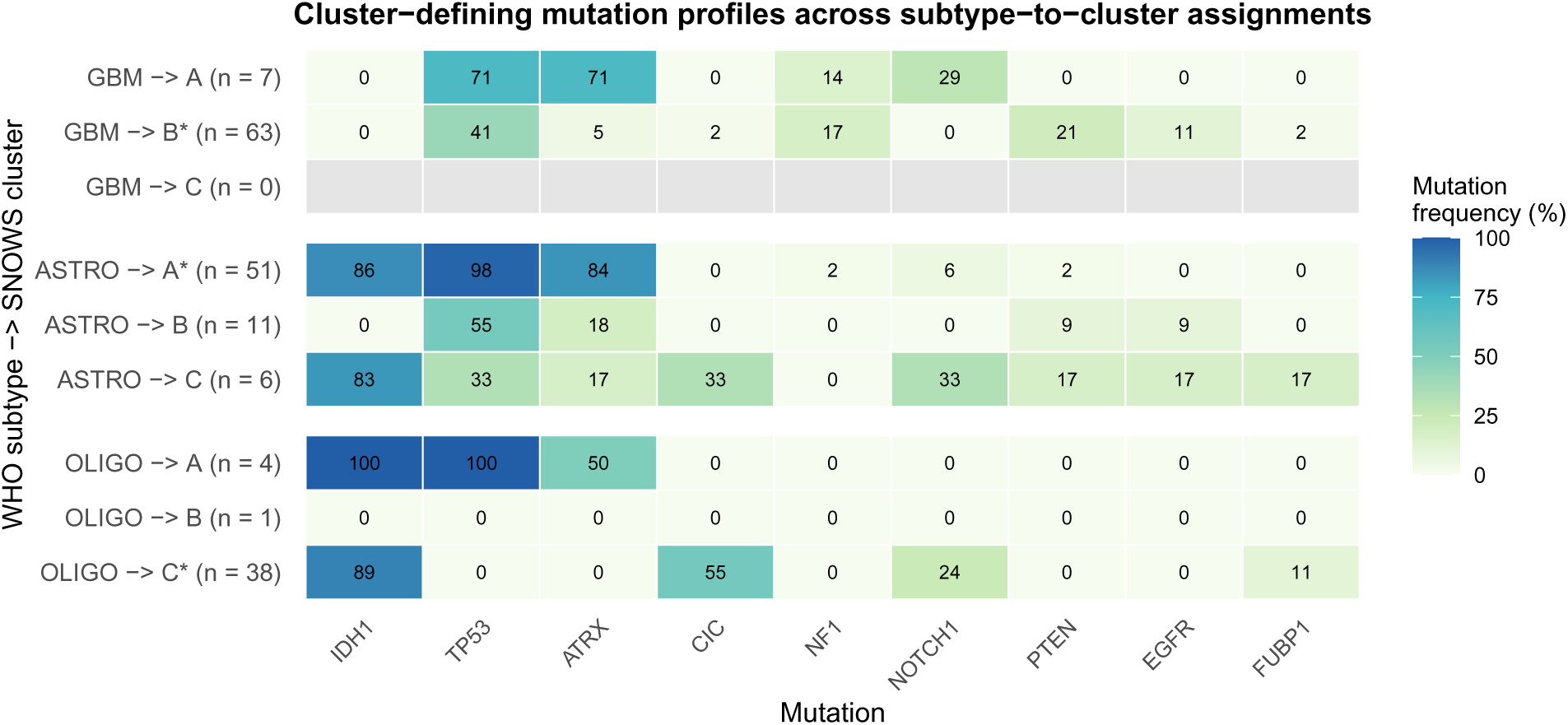
Mutation frequencies across WHO subtype–cluster combinations in CGGA. Rows stratify patients by recorded WHO 2021 subtype and assigned SNOWS cluster, and columns show genes significantly enriched or depleted in the cluster-level analysis. Cell values are mutation frequencies (%). Asterisks identify the cluster containing the largest number of patients of the indicated WHO subtype. Grey cells denote combinations with no patients.

**Table 8:** Prognostic benchmarking of SNOWS and alternative clustering methods in CGGA. Clinical-only models included WHO 2021 subtype, age, and biological sex; full models additionally included cluster assignment. Incremental metrics compare the full and clinical-only models. Dashes indicate IBS estimates that could not be calculated because the corresponding full Cox model contained non-estimable coefficients due to redundant predictors.

| Metric | SNOWS | CANclust | BNMM | k-means |
|---|---|---|---|---|
| Log-likelihood gain (Δℓ) | 7.6 | 6.0 | 0.4 | 1.4 |
| Likelihood-ratio test *p*-value | 4.9 × 10<sup>−4</sup> | 2.4 × 10<sup>−3</sup> | 0.667 | 0.241 |
| C-index, clinical-only model | 0.691 | 0.691 | 0.691 | 0.691 |
| C-index, cluster-only model | 0.700 | 0.691 | 0.670 | 0.695 |
| C-index, subtype-only model | 0.690 | 0.690 | 0.690 | 0.690 |
| C-index, full model | 0.712 | 0.707 | 0.693 | 0.701 |
| ΔC-index | 0.021 | 0.016 | 0.002 | 0.010 |
| Paired concordance test *p*-value | 3.4 × 10<sup>−12</sup> | 1.7 × 10<sup>−9</sup> | 0.0187 | 0.0469 |
| IBS, clinical-only model | 0.170 | 0.170 | 0.170 | 0.170 |
| IBS, cluster-only model | 0.157 | 0.160 | 0.188 | 0.173 |
| IBS, subtype-only model | 0.171 | 0.171 | 0.171 | 0.171 |
| IBS, full model | 0.156 | 0.160 | 0.170 | – |
| ΔIBS | -0.014 | -0.010 | 0.001 | – |
| McFadden pseudo-*R*<sup>2</sup> | 0.561 | 0.528 | 0.773 | 0.836 |
| Cluster novelty index | 0.439 | 0.472 | 0.227 | 0.164 |
| NMI with WHO 2021 subtype | 0.537 | 0.500 | 0.766 | 0.834 |
| NMI with sex | 0.016 | 0.014 | 0.005 | 0.004 |
| NMI with age | 0.089 | 0.101 | 0.125 | 0.129 |
| Cluster size range | 44–75 | 44–71 | 48–71 | 49–70 |
| Number of patients | 181 | 181 | 181 | 181 |

#### A.6.2 Additional cancer cohorts

For the pan-kidney and pan-myeloid cohorts, the number of clusters was selected using BIC on the complete cohort, without survival information, yielding *K* = 3 and *K* = 6, respectively. The selected *K* was fixed throughout five-fold outer cross-validation, with the survival weight *α* tuned by inner cross-validation within each outer training set. The pseudocount was fixed to the glioma-selected value (*χ* = 0.001). Models were refitted on each outer training set and applied to the corresponding held-out fold. Results are reported as means across the five outer folds (Supplementary Table 9).

**Table 9:**
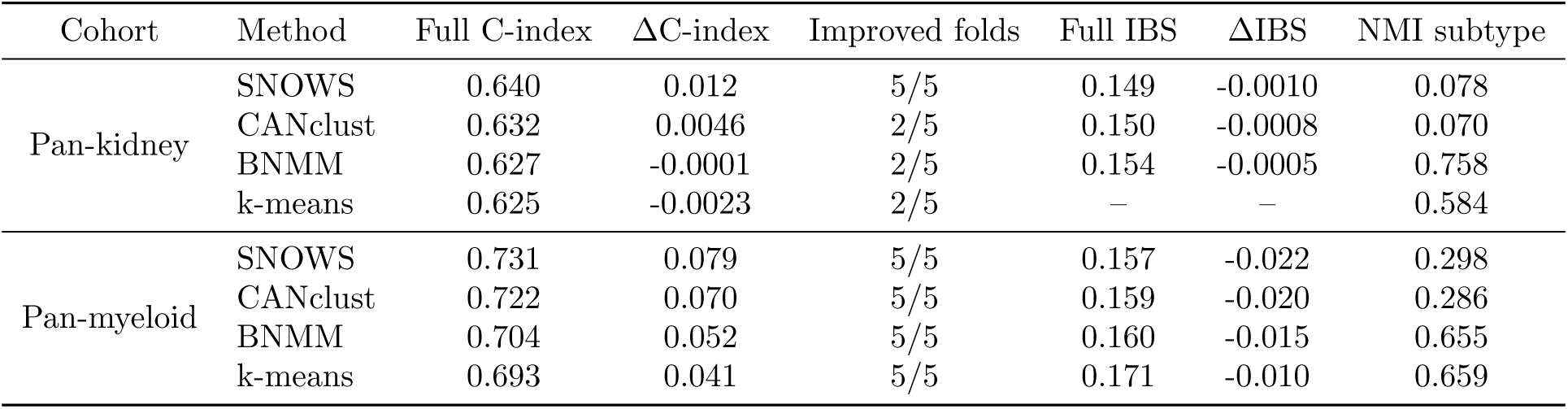
Nested cross-validation results in pan-kidney cancer and pan-myeloid malignancies. Values are means across held-out outer folds. “Improved folds” is the number of outer folds in which the full model had a higher C-index than the clinical-only model. Dashes indicate IBS estimates that could not be calculated because the corresponding full Cox model contained non-estimable coefficients due to redundant predictors.

## Notes

https://github.com/sysbiomed/SNOWS

### Competing Interest Statement

The authors have declared no competing interest.

